# Chronic activation of Toll-like receptor 2 induces an ichthyotic skin phenotype

**DOI:** 10.1101/2022.06.06.494922

**Authors:** Hephzi Tagoe, Sakinah Hassan, Gehad Youssef, Wendy Heywood, Kevin Mills, John I. Harper, Ryan F.L. O’Shaughnessy

**Affiliations:** Centre for Cell Biology and Cutaneous Research, Queen Mary University of London, UCL Great Ormond Street Institute of Child Health, UK; Livingstone Skin Research Centre, UCL Great Ormond Street Institute of Child Health, UK; Immunobiology, UCL Great Ormond Street Institute of Child Health, UK; Dermatology and Genetics and Genomic Medicine, UCL Great Ormond Street Institute of Child Health, UK

**Author notes:** **Corresponding author:** Ryan O’Shaughnessy PhD, Centre for Cell Biology and Cutaneous Research, Blizard Institute, Queen Mary University of London, 4 Newark 4 Newark Street, London E1 2AT. **Funding sources:** British Skin Foundation, Foundation for ichthyosis and related skin types, Great Ormond Street Children’s Charity.

**Keywords:** Hyperkeratosis, Toll-Like receptor, Gata3, Ichthyosis

## Abstract

Ichthyosis defines a group of chronic conditions that manifest phenotypically as a thick layer of fish-like scales in response to disorders of cornification and often affects the entire skin. While the gene mutations that lead to ichthyosis are well documented, the actual signalling mechanisms that lead to scaling are poorly characterised, however recent publications suggest that there are common mechanisms active in ichthyotic tissue, and in analogous models of ichthyosis. Combining gene expression analysis of gene-specific shRNA knockdowns of more severe autosomal recessive congenital ichthyoses (ARCI) and proteomic analysis of skin scale from ARCI patients, we identified a common activation of the Toll-like receptor (TLR) 2 pathway. Exogenous activation of TLR2 led to increased expression of important cornified envelope genes and in organotypic culture caused hyperkeratosis. Conversely blockade of TLR2 signalling in ichthyosis patient keratinocytes and our shRNA models reduced the expression of keratin 1, a structural protein overexpressed in ichthyosis scale. A time-course of Tlr2 activation in rat epidermal keratinocytes revealed that although there was rapid initial activation of innate immune pathways, this was rapidly superseded by widespread up-regulation of epidermal differentiation related proteins. Both NFκβ phosphorylation and the Gata3 up-regulation was associated with this switch and Gata3 overexpression was sufficient to increase Keratin 1 expression. Taken together, these data define a dual role for Tlr2 during epidermal barrier repair, that may be a useful therapeutic modality in treating diseases of epidermal barrier dysfunction.

## Introduction

The ichthyoses are a family of mendelian diseases of keratinisation (Oji et al., 2010) in which the cornified envelope - the principal barrier-conferring component of the epidermis - becomes defective in either the lipid or protein component (Takeichi et al., 2016). Patients with autosomal recessive congenital ichthyosis (ARCI) are the most severely affected. Most children with ARCI present as a collodion baby at birth (Dyer et al., 2013), but shortly after thick scales appear in most cases over the entire skin of the patient. Due to improvements in neonatal care, there is a minimal risk of perinatal death due to insensible water loss and infections, however their skin remains abnormal throughout life, and is the principal phenotype in ARCI.

ARCI can be subdividing into subclasses of disease defined by diagnostic criteria. Lamellar ichthyosis (LI) typically presents at birth with a collodion, which is shed, after which thick scaling is present without redness of the skin, while non-bullous ichthyosiform erythroderma (NBIE) has lighter scaling, but also presents with redness of skin, and collodion is less common, but still occurs (Oji et al., 2010; Vahlquist et al., 2010). Approximately 90% of LI is caused by mutations in the transglutaminase 1 gene (TGM1) (Russell et al., 1995), while TGM1 mutations are only rarely associated with NBIE (Vahlquist et al., 2010) Instead, mutations in the arachidonate lipoxygenases ALOX12B and ALOXE3 predominate (Hotz et al., 2021; Jobard et al., 2002). Therefore, although there are many genes now identified as being mutated in ARCI (Uitto et al., 2021), they all encode genes important to epidermal barrier function.

The scaling typically seen in lamellar ichthyosis and non-bullous ichthyosiform erythrodrma reflects a homeostatic response to the chronic barrier dysfunction. Grafting Tgm1 and Alox12b knockout skin on immunocompromised mice revealed that barrier function was restored through hyperkeratosis and increased lipid production (Kuramoto et al., 2002; de Juanes et al., 2009). Understanding the genes expression changes underlying this hyperkeratotic response will be critical in finding therapies to reduce scaling in these patients. Oral retinoids are the primary treatment for these patients, and they efficiently reduce patient scale, but they are associated with serious side effects (Rood et al., 2007). Although gene-specific gene therapy and protein replacement therapies are being developed (Choate et al., 1996; Aufenvenne et al., 2013; Plank et al., 2019), mechanism-specific, generalised therapies to reduce scaling are still greatly needed. Examining the effects of Tgm1 loss versus Alox12b loss in keratinocytes has the potential to define differences between these two disorders, while looking at common changes between these two knockdown lines defines a common pathway to hyperkeratosis and scaling in ARCI (Youssef et al., 2013)

Over the last twelve years, significant in-roads have been made in understanding the mechanisms underlying scaling in ARCI, In Tgm1 and Alox12b shRNA knockdown keratinocytes, we have identified IL1A signalling and activation and upregulation of IL1A and MDM2 as necessary for hyperkeratosis in our 3D culture models (O’Shaughnessy et al., 2010; Youssef et al., 2013). MDM2 upregulation increased the expression of the key structural keratin, keratin 1, but not keratin 10 (Youssef et al., 2013). Analysis of full thickness skin biopsies from ichthyosis patients indicated a Th17/IL23/IL-36-related signature of upregulated genes in both the dermis and epidermis of patients (Malik et al., 2019; Paller et al., 2017), and in some cases up-regulation of these signature genes correlated with increased trans-epidermal water loss (Brunner et al., 2018). ABCA12 shRNA knockdown lines, which phenocopied the severe ichthyosis, harlequin ichthyosis, also showed upregulation of key innate immune mediators, such as STAT1, and IL-36 family cytokines and NOS2 (Enjalbert et al., 2020). Taken together, this strongly implicates activation of innate immune pathways in ARCI.

A greater understanding of the key keratinocyte-mediated molecular mechanisms underlying altered innate immune signalling and resultant scaling in these patients would facilitate the identification of new therapeutic targets and the subsequent discovery of molecular modulators of scaling. In this study, we identified novel upregulated pathways in ARCI using an integrated analysis of proteomics data from patient scales and our datasets from our ARCI *in vitro* models. We identified upregulation of the toll-like receptor 2 pathway. Endogenous activation of TLR2 caused scaling in organotypic models. Blocking TLR2 signalling in our ichthyosis models reduced markers of hyperkeratosis, such as keratin 1. RNAseq analysis of a time course of TLR2 activation in keratinocytes induces a biphasic gene expression pattern, a short term innate immune response, and a longer term, GATA3-mediated hyperkeratotic response. These changes mirror the effects seen in Alox12b knockdown and NBIE patient keratinocytes.

## Materials and Methods

### QTOF mass spectrometry

QTOF mass spectrometry and subsequent analysis were described previously (Bennett et al., 2010). Peptides were analysed using a nanoAcquity ultrahigh definition liquid chromatography (UPLC) system coupled to a quadrupole time-of-flight (QToF) Premier mass spectrometer (MS) (Waters Corporation, Manchester, UK). Peptides were trapped and desalted before reverse phase separation using a 5 mm x 300 μm Symmetry C18 5 μL, pre-column. Peptides were separated using a 15 cm x 75 μm C18 reverse phase analytical column and loaded onto the pre-column in a 3% acetonitrile (ACN) and 0.1% formic acid (FA) in ultrapure water solution (Fluka) at a flow rate of 4 μL/min for 4 min. Peptides were eluted from and separated on the analytical column using a 3-40% ACN gradient containing 0.1% FA in ultrapure water over 30 min at a flow rate of 0.3 μL/min. The column was re-equilibrated to the starting conditions for 9 min, after removal of the non-polar and non-peptide material with 100% ACN containing 0.1% FA for 5 min at a flow rate of 0.4 μL/min. Columns were maintained at 35 °C and mass accuracy was maintained during the run using 0.3 nmol/L of [glutamic acid1]-fibrinopeptide B delivery through an auxiliary pump of the nanoAcquity at a flow rate of 0.3 μL/min57.

Peptides were analysed in positive ion mode using a QToF Premier (Waters Corporation, Manchester, UK), operated in V-mode, with a typical resolving power of 10000 fwhm. The ToF analyser was calibrated prior to analyses with [glutamic acid1]-fibrinopeptide B fragments over the mass range of 50-2000 m/z obtained using 25 eV of collision energy. Data files were mass-corrected every 30 s using the doubly charged [glutamic acid1]-fibrinopeptide B species (785.84262 m/z). Accurate mass LC-MS data were collected in a data independent and alternating, low and high collision energy mode. Each low/high acquisition was 1.5 s in duration with a 0.1 s inter-scan delay. Low energy data collections were performed at constant collision energy of 4 eV, high collision energy acquisitions were performed across a 15−40 eV ramp over 1.5 s and a complete low/high energy acquisition was achieved every 3.2 s.

Raw data were imported into Nonlinear Dynamics Progenesis QI for proteomics to identify peptide masses corresponding to the fragmentation ion data. Mass correction was based on [glutamic acid1]-fibrinopeptide B delivered via an auxiliary pump. Processed spectra were merged prior to searching the UniProt reviewed human proteome. Search parameters were set to two fragment ions matched per peptide, four fragment ions per protein and two peptides per protein and one missed cleavage, fixed modifications were carbamidomethylation of cysteines and dynamic modifications were hydroxylation of aspartic acid, lysine, asparagine and proline and oxidation of methionine and false discovery rate was 4 %.

### Cell culture

Rat epidermal keratinocytes (REKs) were used in some of these experiments, as because they are spontaneously transformed, they grow readily in culture without the use of a feeder layer but they can still fully differentiate without the use of calcium switch (Baden and Kubilus, 1983). REKs of passages between 20 and 30 were passaged in DMEM +10% fetal bovine serum and incubated at 37°C and 5% CO2. For immunofluorescence analysis, the cells were subsequently simultaneously fixed and permeabilized in 4% paraformaldehyde/0.2% Triton X-100. Organotypic culture was performed as described by O’Shaughnessy et al. (O’Shaughnessy et al., 2007), briefly, 2×10^5^ rat epidermal keratinocytes were cultured in medium containing 100 mg/ml G418 on de-epidermized dermis made from cadaverous skin (Euro Skin Bank, Beverwijk, Netherlands) in a metal ring until confluent. Subsequently the organotypic cultures were raised to the air-liquid interface and cultured for a further 10 days, unless specified. organotypics were incubated at 37°C and 10% CO_2_. The constructs were processed for paraffin embedding by fixing in 4% paraformaldehyde.

Human epidermal keratinocytes (HEKs, Thermofisher), and patient keratinocytes, passages less than 5, were cultured in Defined Epilife supplemented medium. This was prepared by adding 5ml of Epilife defined growth supplement to the base medium (Thermofisher). Keratinocytes were then differentiated by gradual calcium switch from 0.06mM to 2.4mM CaCl_2_. (Bilke et al, 2012). For the inhibitor and drug treatments we used the following drugs at the following concentrations. PAM3CSK4 (Tocris bioscience) 10nM, CU-CPT-22 (Sigma Aldrich) 0.4 mM. Etoposide (7.5μg/ml, Sigma, UK), LPS (100nL/ml. Sigma, UK), DTT (2.5μl/ml, Invitrogen, UK), and Tunicamycin (50ng/ml, Sigma, UK)

Alox12b and Tgm1 shRNA knockdown in rat epidermal keratinocytes using SureSilencing shRNA plasmids (Qiagen) has been previously described (O’Shaughnessy et al, 2010; Youssef et al., 2013). The GATA3 construct for overexpression in rat epidermal keratinocytes was obtained from OriGene. All transfections were performed using lipofectamine 2000 (ThermoFisher), according to manufacturers’ instructions.

### Immunofluorescence

Keratinocytes were fixed in 4% paraformaldehyde containing 0.2% Triton-X-100 before blocking in PBS containing 0.2% Triton-X-100 and 0.2% Fish skin Gelatin. Primary and secondary antibodies were also incubated in this blocking medium. Slides were mounted with DAPI-containing Prolong gold (Thermofisher). Images were taken using a Leica DM4000 upright epifluorescence microscope. The following antibodies were used at the following concentrations, Rabbit anti-pNFDB (pp65) (3033s, Cell Signalling technology), 1:50; Rabbit anti-Sumo 1 (4940, Cell Signalling Technology) 1:100; Rabbit anti-Sumo 2/3 (4971, Cell Signalling Technology) 1:100.

### Western blotting

Proteins were extracted in total lysis buffer 10% SDS, 10%, β-mercaptoethanol 10mM Tris at pH7.5, and run on 10-20% TGX polyacrylamide gel (Bio-Rad) and blotted onto Hybond N nitrocellulose filters (Amersham). Nonspecific binding sites on the membrane were blocked by incubating the membrane in 5% milk powder in PBS containing 1X Tween-20 (Sigma). The membrane was then incubated with the primary antibody of interest at 4^0^C overnight. A secondary antibody conjugated to horse radish peroxidase was incubated for 60mins at room temperature. The membrane was developed in ECL chemiluminescent western blotting detection reagent (Amersham) and bands were detected on film (Amersham). The following antibodies were used. Rabbit anti-GATA3 (10417-1-AP, Proteintech), 1:500; Rabbit anti-Keratin 1 (905601, Biolegend), 1:2000; Rabbit anti-Keratin 10 (905401, Biolegend), 1:5000; Rabbit-anti IDβα (9242s, Cell Signalling Technology); 1:500. Rabbit anti-pNF-DB (pp65) (3033s, Cell Signalling technology), 1:250; Rabbit anti-NF_\ll_β (p65) (4764s, Cell signalling); Mouse anti-GAPDH (MAB374, Millipore) 1:2000; Rabbit anti-IL1a (sc-7929, Santa Cruz Biotechnology) 1:2000; Rabbit anti-Loricrin (905101, Biolegend) 1:1000; Mouse anti-Involucrin (GTX42415, Genetex) 1:1000; Rabbit anti-TLR2 (NB100-56573, Novus Biologicals) 1:500. Rabbit anti-Sumo 1,2,3 (A-714, Biotechne) 1:500; Rabbit anti-pSer71 Rac1 (2461, Cell Signalling technologies) 1:500.

### RNAseq analysis and downstream analyses

#### REKs were treated with PAM3CSK4

4 at 2, 6, 12, and 24 hours followed by RNA extraction and purification using the RNeasy kit (Qiagen). Samples were checked for purity and concentration using a nanodrop and Bioanalyser. Libraries were prepared using NEB mRNA library prep kit from 100ng total RNA as per manufacturer’s instructions. Samples were subsequently checked for quality using Quibit and Tape station. Sequencing was done on a single lane of Illumina NextSeq 500 using a high output kit 75bp paired end reads. Principal Component Analysis (PCA) on the gene intensity data was run on the gene expression data followed by a two-way ANOVA to determine differentially expressed genes across all timepoints. Heat maps derived from these data and previous data (Youssef et al., 2013) were made using Morpheus (https://software.broadinstitute.org/morpheus). STRING analysis (https://string_dg.org) was performed using the full string network of both functional and physical associations, with a minimum confidence of 0.400 for positive evidence of association. Gene ontology analyses were performed using DAVID (https://david.ncifcrf.gov; Huang et al., 2009a; Huang et al., 2009b), Mapping of rat gene expression data to human promoter binding was performed with Enrichr (https://maayanlab.cloud/Enrichr/) (Chen et al., 2013)

### Statistical and image analysis

All data analysed is based on mean values. Protein expression by SDS-PAGE and western blot has been analysed in imageJ software using densitometry with a limit to the band threshold. This was followed by calculating the fold change to untreated control using normalised densitometry values in excel. T-test was used to assess the statistical significance between the treated and untreated samples as well as analysis of variance (ANOVA), followed by post-hoc testing. Gene expression changes were analysed using 2 – tailed -t-test. RNAseq data was analysed by principal component analysis and two-way ANOVA, and cell staining intensity has been quantified in image J software. Error bars on biological repeats show the standard deviation of the mean.

## Results

### Combining proteomic and gene expression data revealed an upregulated SUMO1/TLR2/GATA3 axis in ARCI

To identify changes in expression in innate immune pathways we combined our previously determined genes that were commonly upregulated in Tgm1 and Alox12b knockdown rat epidermal keratinocytes (REKs) (O’Shaughnessy et al., 2010; Youssef et al, 2013), a spontaneously transformed rat keratinocyte line that is able to fully differentiate in culture (Baden and Kubilus, 1983). with a shotgun proteomic analysis comparing scale from 4 individuals with ARCI, and 8 normal control scale samples. We identified 20 proteins upregulated in and 18 proteins downregulated by at least 1.5-fold in two or more patient samples (Figure 1A and B; Supplementary table ST1). Of these, the most highly upregulated proteins were Cathepsin D (CTSD), APOA1 and the Sumo ligase RANBP2. Gene ontology analysis revealed over-representation of proteins involved in the formation of the cornified envelope and keratinisation, sumoylation and toll-like receptor (TLR) signalling (Figure 1C). As only a proportion of proteins are typically detectable by label free mass spectrometry of cornified envelopes (Karim et al., 2018; Winget et al., 2016; Bennett et al., 2010), we combined this data set with our previous gene expression analysis of Tgm1 and Alox12b shRNA knockdowns and identified a large, interrelated protein-protein interaction network with a potential functional connection between RANBP2, SUMO1, and GATA3 (Figure 1D). We confirmed overexpression of Sumo1, but not Sumo2/3 in our ARCI models in the array hybridisation data (Figure 1E), which was confirmed by immunofluorescence and (Figure 1F and G). Additionally, by western blot using a pan SUMO antibody, we identified an increase in sumoylated protein species in our ARCI shRNA knockdown models (Figure 1H).

**Figure 1.**
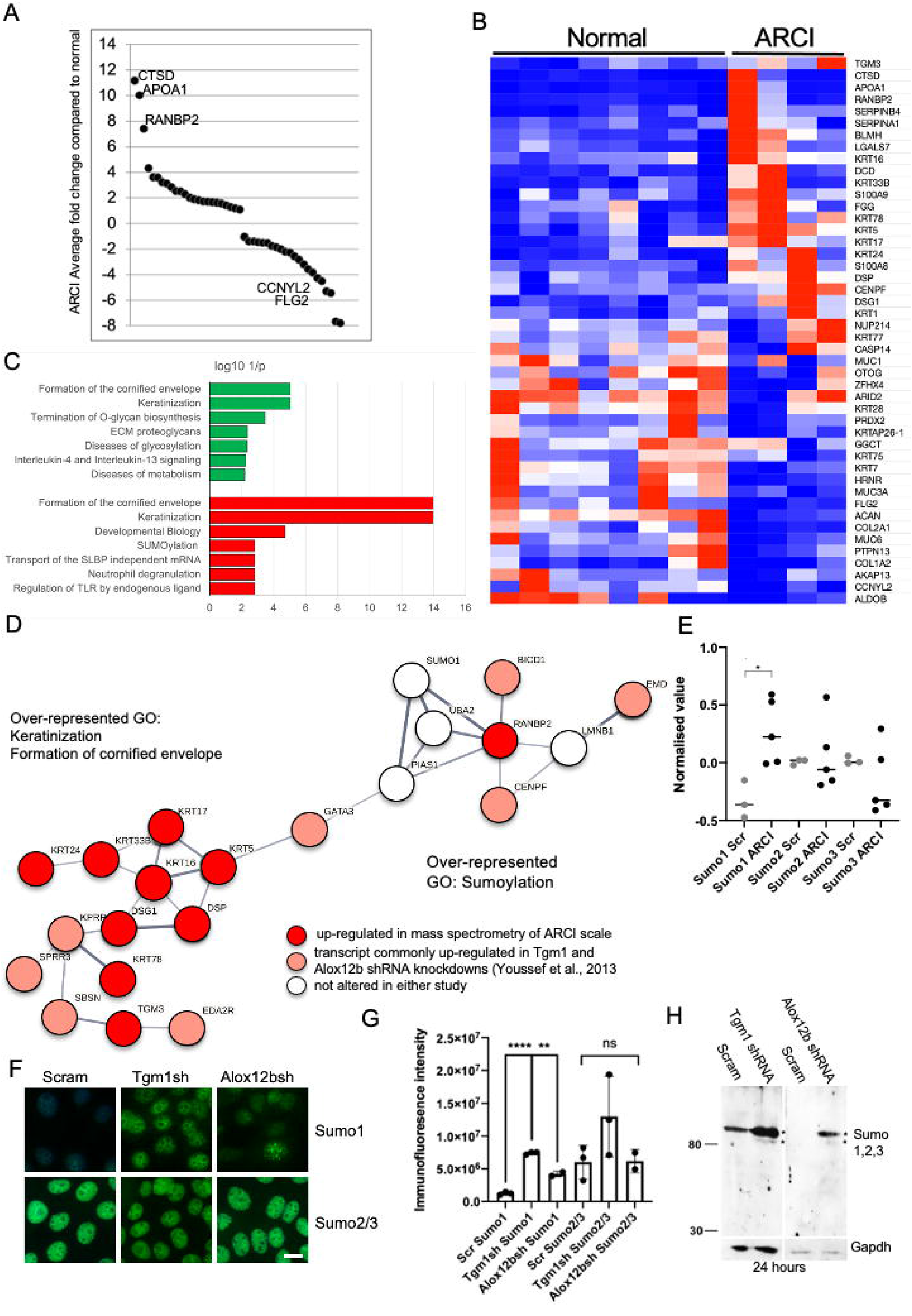
Combining proteomic and gene expression data revealed an upregulated SUMO1/TLR2/GATA3 axis in ARCI. A) Graph of average fold-change of protein expression from the protein mass spectrometry data from ARCI patient scale compared to normal. B) Heat map of these data, red is maximum values and blue is the minimum value on each row. C) bar graph showing over-represented biological process gene ontologies in the up-regulated and down-regulated proteins (red and green respectively). D) String network analysis of combined over expressed proteins in scale and genes that are differentially expressed in both Tgm1 and Alox12b shRNA knockdown rat epidermal keratinocytes (Youssef et al., 2013). E) Graph of expression of each of the SUMO isoforms in control rat epidermal keratinocytes (Scr) and in both the Tgm1 and Alox12b shRNA knockdowns (ARCI), p<0.05 T-Test. F) Immunofluoresence of Sumo 1 and Sumo 2/3 in control rat epidermal keratinocytes (Scr) and in both the Tgm1 and Alox12b shRNA knockdowns, n=3, bar 10 µm. G) Graph of Sumo1 and Sumo2/3 immunofluorescence intensity,**p<0.005, ****p<0.00005 One way ANOVA follows by post-hoc testing H) Western blot of a pan-SUMO antibody in control rat epidermal keratinocytes (Scr) and in both the Tgm1 and Alox12b shRNA knockdowns. * indicates over-represented sumoylated proteins species.

### Activation of TLR2 induced hyperkeratosis in organotypic culture

Based on the potential functional interaction between SUMO1 and TLR2, we tested the hypothesis that TLR2 induction could activate SUMO expression in keratinocytes by treating with either Lipopolysaccharide (LPS), which activates TLR4 or the synthetic ligand PAM3CSK4 which activates TLR2 (Figure 2A) for 24 hours. By immunofluorescence we observed Sumo1 upregulation only after PAM3CSK4 treatment. Testing a number of different stressors (Supplementary Figure S1A), indicated that Sumo1 up-regulation in REKs was only observed in PAM3CSK4 treatment. By western blot we saw significantly increased expression of keratin 1, without con-comitant expression of keratin 10, and upregulation of IL1A only in cells treated with LPS or PAM3CSK4. Other stressors, the DNA synthesis inhibitor etoposide or the ER stress inducer tunicamycin, failed to induce either Keratin 1 or Il1A (Supplementary figure S1B and C)

**Figure 2.**
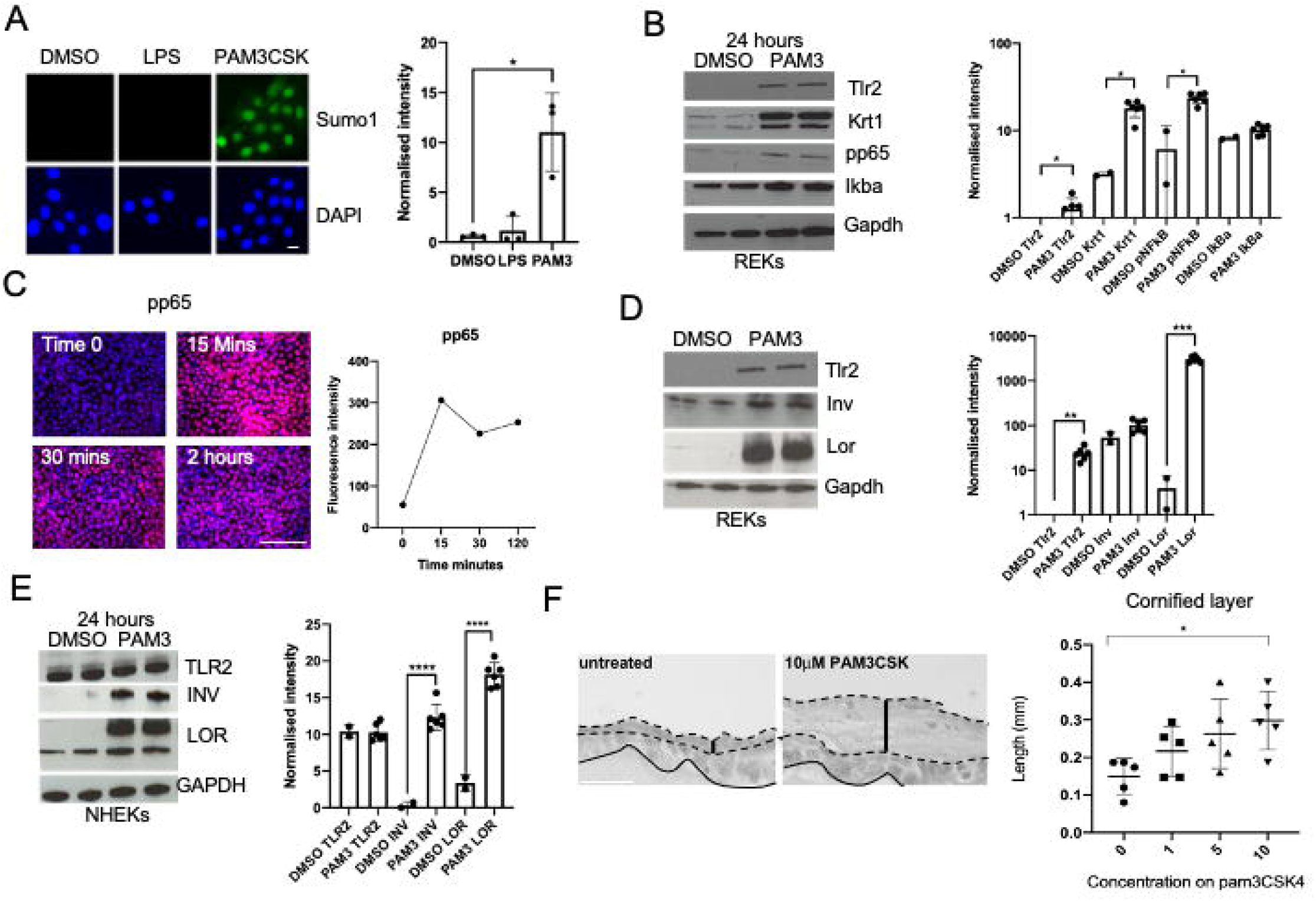
Activation of TLR2 induced hyperkeratosis. A) Sumo 1 immunofluoresence in rat epidermal keratinocytes treated with LP, PAM3CSK4, or vehicle (DMSO). DAPI is a nuclear counterstain. Graph shows normalised intensity (n=3) B) Western blot of Tlr2, keratin 1 (Krt1), phosphor-p65 (pp65) and Ikba. In rat epidermal keratinocytes treated with PAM3CSK4 (PAM3, n=6) or vehicle (DMSO, n=2). Gapdh serves as loading control. Graph shows normalised intensity. D) pp65 immunofluoresence in rat epidermal keratinocytes over a 2 hour timecourse with intensity graph on the right. D and E) Western blots for Toll-like receptor 2 (Tlr2/TLR2), Involucrin (Inv/INV) and Loricrin (Lor/LOR) in PAM3CSK4 (PAM3, n=6) or vehicle (DMSO, n=2) treated rat epidermal keratinocytes (D) and Normal human epidermal keratinocytes (E). Graphs show normalised intensity F) Histology of rat epidermal keratinocytes either untreated or treated with 10mm PAM3CSK4 from 5 days post raising to the air-liquid interface until day 10. Continuous line indicates the dermo-epidermal junction and the dotted lines indicate the extent of the cornified layer. The thick bar indicates the thickness of the cornified layer. F) Quantification of cornified layer thickness in PAM3CSK4 treated organotypic cultures. I measured at n=5 sites over 2 different organotypic cultures in ImageJ. *, p <0.05, ** p<0.005, ***p<0.0005, p<0.00005, one-way ANOVA followed by post-hoc testing, ns = not significant. Error bars, SD, Bar 10µm (A), 50µm (D and G)

We also checked that the Tlr2 signalling pathway was up-regulated in our Tgm2 and Alox12b shRNA knockdown rat epidermal keratinocytes. While Tlr2 mRNA expression was not significantly increased (Supplementary figure S2), Tlr2 protein expression was increased in the shRNA knockdown lines (Figure 2B). There was also increased phosphorylation of the p65 subunit of NFκβ, consistent with our previous finding of nuclear localisation of NFκβ in Tgm1 knockdown organotypic cultures (O’Shaughnessy et al., 2010). However, Tlr2 activation by PAM3CSK4 caused p65 phosphorylation without Iκβα degradation (Figure 2B). Despite this, Nuclear localisation of pp65 still occurred within 15 minutes of PAM3CSK4 treatment and was maintained for 2 hours (Figure 2C). In both REKs and NHEKs, 24 hours of PAM3CSK4 treatment upregulated key epidermal terminal differentiation markers keratin 1 (Figure 2C), loricrin, and involucrin (Figure 2D and E). However it was notable that while in REKs this was associated with an increase in Tlr2 protein, in NHEKs, there was significant TLR2 protein present before PAM3CSK4 treatment and no change after treatment. In skin equivalent organotypic culture, PAM3CSK4 treatment induced a dose-dependent thickening of the cornified layer, without increased vital epidermal thickness (Figure 2F).

### TLR2 inhibition reduced levels of Keratin 1 in both ARCI models and in ARCI keratinocytes

CU-CPT22 is a selective inhibitor of TLR2 (Cheng et al., 2012). CU-CPT22 treatment of Tgm1 and Alox12b shRNA expressing keratinocytes reduced keratin 1 expression over an 8-hour period (Figure 3A), with a longer-lasting effect observed in Alox12b shRNA knockdowns. We also treated non-genotyped ARCI primary keratinocytes from 2 patients diagnosed with lamellar ichthyosis (LI) and non-bullous ichthyosiform erythroderma (NBIE) both showed increased keratin 1 expression compared to normal keratinocytes, while IL1A was increased only in LI keratinocytes (Figure 3B). CU-CPT22 treated caused a transient reduction of keratin 1 expression after a single treatment (Figure 3C). Taken together these data suggest that activation of the TLR2 pathway is required for hyperkeratosis in ichthyosis, and that inhibition of this pathway can reduce the expression of keratin 1, a key marker of hyperkeratosis.

**Figure 3.**
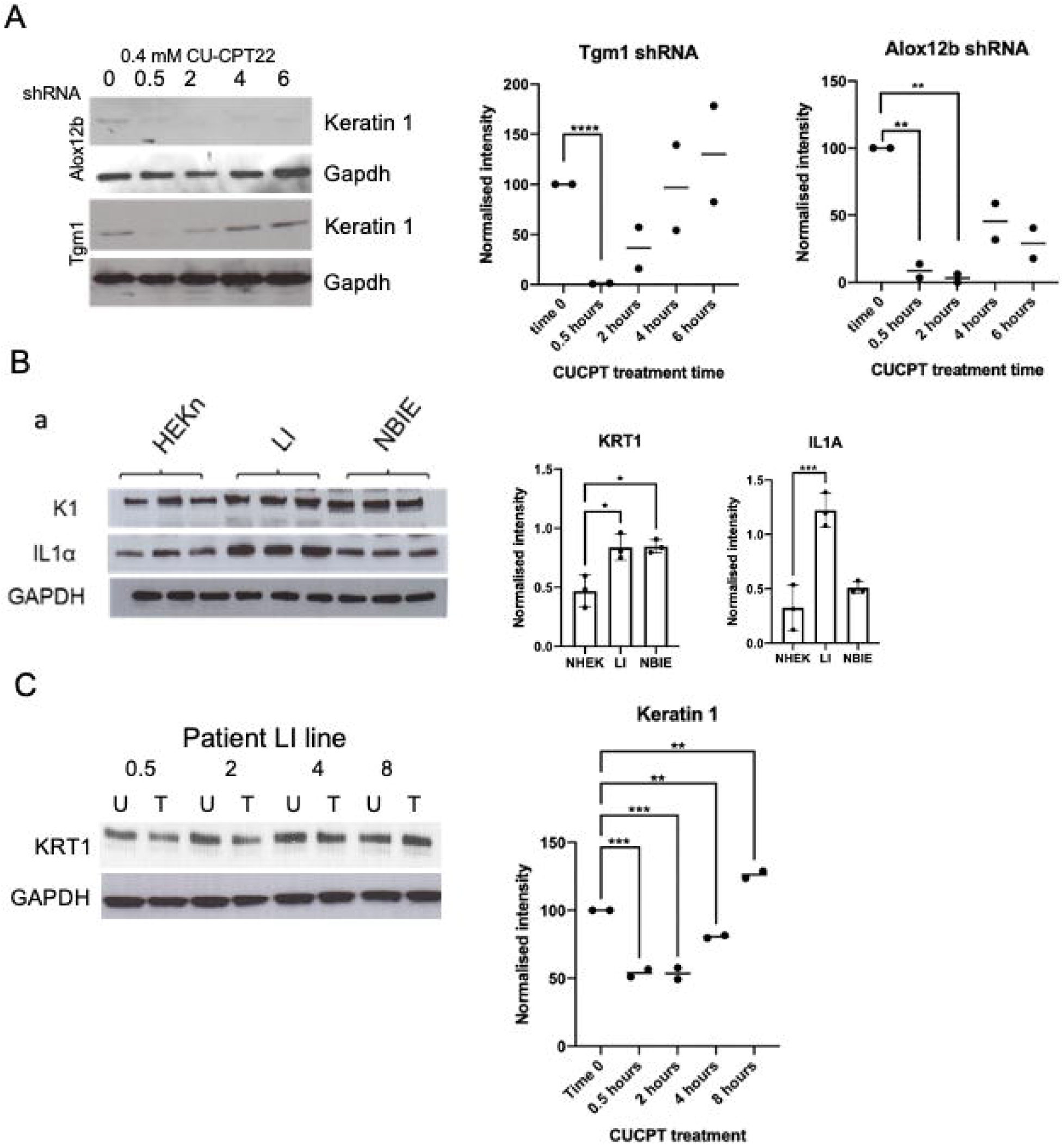
TLR2 inhibition reduced levels of Keratin 1 in both ARCI models and in ARCI keratinocytes. A) Western blot of keratin 1 in Tgm1 and Alox12b shRNA expressing keratinocytes treated with the TLR2 inhibitor CU-CPT22 over an 8 hour timecourse. Gapdh serves as a loading control. Graphs show densitometry of n=2 biological replicates, Alox12b and Tgm1 shRNA expressing rat epidermal keratinocytes treated with CU-CPT22 showing comparison as a % of the untreated time zero value. B) Triplicate westerns for keratin 1 (K1) and interleukin 1a (IL1a) in cell lines from 2 ARCi patient, one with lamellar ichthyosis (LI), and one with non-bullous ichthyosiform erythroderma (NBIE), not sequenced. GAPDH serves as loading control. Graphs show normalised densitometry of keratin 1 and IL1A in both patient lines. C) Representative western blot (n=2) of keratin 1 (KRT1) in the lamellar ichthyosis patient line treated once with CU-CPT22 at time 0 over an 8-hour timecourse. U, untreated, T, treated. H) Graph of normalised intensity of western blot. *, p <0.05, ** p<0.005, ***p<0.0005, ****p<0.00005, one-way ANOVA with post-hoc testing.

### RNAseq analysis and signalling pathway analysis of rat epidermal keratinocytes treated with PAM3CSK4 indicated a biphasic pattern of NFκβ activation

After a single treatment with PAM3CSK4 there was an increase of keratin 1 and IL1A after 12 hours, and pp65 showed both early and late increases in phosphorylation (Figure 4A). This suggested that multiple phases of gene expression changes were occurring culminating in increased expression of terminal differentiation markers. To examine this further we performed RNAseq on rat epidermal keratinocytes treated once with 10nM PAM3SCK over a 24-hour period (Figure 4B and C; RPKM expression data in Supplementary Table S4). 1810 genes were differentially expressed over this timescale after applying correction for multiple testing (Figure 4B). Consistent with data from the ARCI shRNA lines, PAM3CSK4 treatment did not significantly increase Tlr2 mRNA expression, although there was a transient increase in Tlr4 at 2 hours and a progressive decrease in Tlr5 expression (Supplementary figure S3)

**Figure 4.**
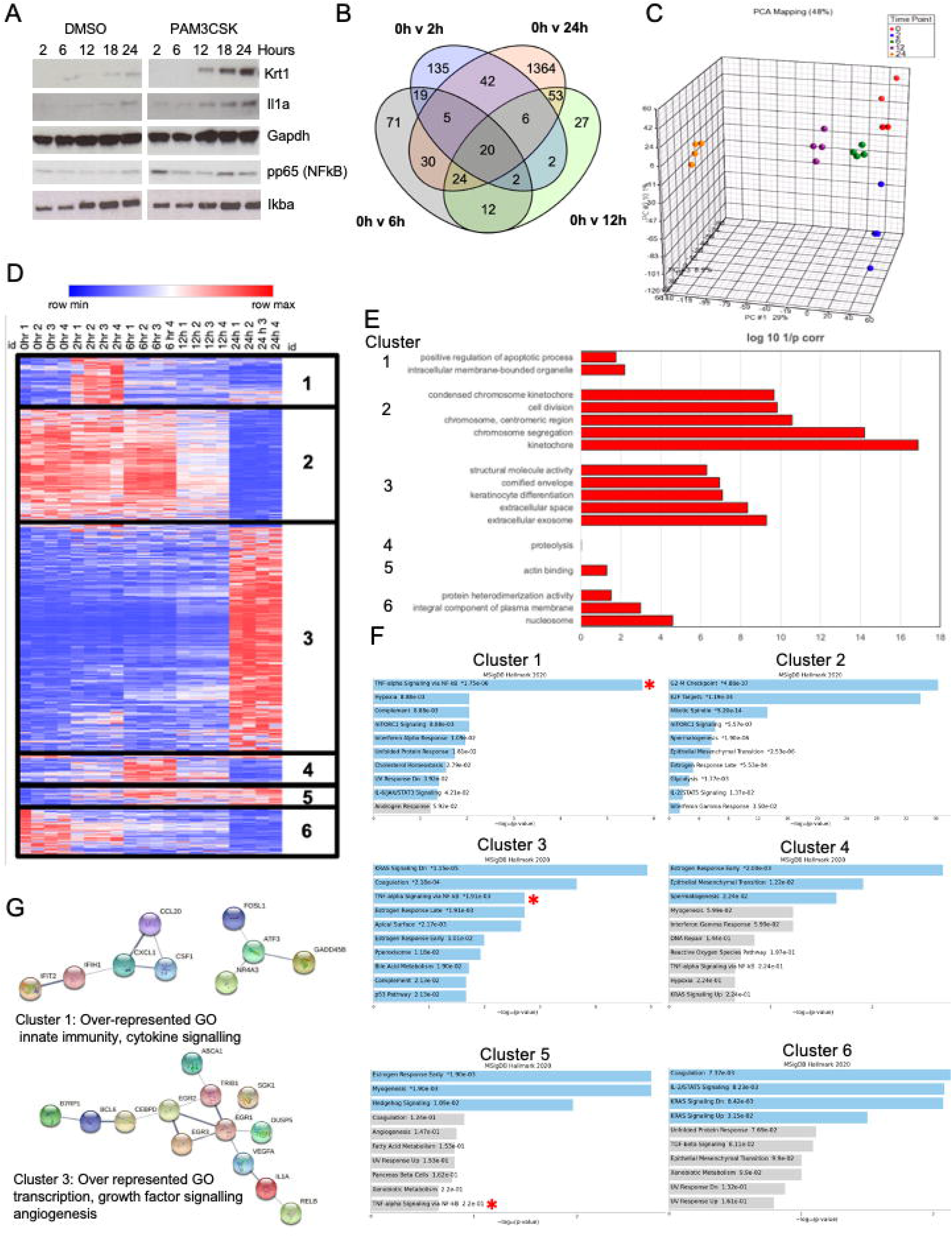
RNAseq analysis and signalling pathway analysis of rat epidermal keratinocytes treated with PAM3CSK4 indicated a biphasic pattern of NFκβ activation. A) Western blot of rat epidermal keratinocytes treated with PAM3CSK4 or vehicle (DMSO) over a 24-hour time-course. GAPDH serves as a loading control. B) Venn diagram showing differentially expressed genes at different timepoints during the 24-hour time-course RNAseq experiment. C) Graphical representation of Principal component analysis showing the variation at both early and late timepoints. D) Heat map and cluster analysis of all differentially expressed genes in the RNAseq experiment. E) Analysis of over-represented gene ontology groups in each of the clusters described in (D). F) Enrichr analysis of signalling related genes in each of the clusters. Blue denotes significant overrepresentation after correction for multiple testing. The red asterisk indicates TNF/NFκβ signalling in the relevant clusters. G) String analysis and gene ontology analysis of the TNF/NFκβ signalling genes in cluster 1 and Cluster 3.

Principal component analysis showed a biphasic change in gene expression. Changes from 0 hours to 2 hours were seen in Principal component 2, while the changes in gene expression at 12 and 24 hours were seen in principal component 1 (Figure 4C). We identified 6 different patterns of gene expression change in our RNAseq data (Figure 4D; Supplementary Table ST2) that reflected the dynamic changes in gene expression, Cluster 1, was transiently overexpressed at the 2 hr timepoint, Cluster 2 was significantly downregulated from 12-24 hours, Cluster 3 were genes maximally expressed at 24 hours. Expression of Cluster 4 genes peaked at 6 hours, expression of Cluster 5 genes increased progressively over the treatment period, Expression of cluster 6 genes fell progressively over the treatment period. Biological function Gene ontology analysis of each cluster (Figure 4E) showed Cluster 1 was associated with positive regulation of apoptosis, Cluster 2 was associated with cell division, Cluster 3 was associated with keratinocyte differentiation and the formation of the cornified envelope and included increased expression of both Keratin 1 and Il1a. Cluster 4 was associated with proteolysis, Cluster 5 was associated with actin binding and Cluster 6 with the nucleosome (Figure 4E).

We analysed the signalling pathways upregulated in each cluster by enrichr (Chen et al., 2013; Figure 4F). TNFa/NFκβ signalling was significantly increased in Cluster 1 and Cluster 3, reflecting the biphasic activation we have already observed. Cluster 2 genes were enriched in genes associated with aspects of the cell cycle, consistent with a switch to increased terminal differentiation. Cluster 4 and 5 were associated with the early estrogen response. Cluster 6 was strongly associated with Ras signalling, again reflecting a switch from proliferation to differentiation. TNFa/NFκβ related genes were different between cluster 1 and 3, with cluster 1 consisting of genes related to innate immunity and cytokine signalling. And cluster 3 consisting of genes involved in transcription, growth factor signalling and angiogenesis (Figure 4G). Taken together these data suggest that NFκβ signalling plays two different roles during the time-course of the response to TLR2 activation.

### Gata3 up-regulation increased keratin 1 expression downstream of TLR2 activation

To determine which of the genes controlled by TLR2 activation are involved in hyperkeratosis in ichthyosis, we compared the gene expression data with our previous analysis of Tgm1 and Alox12b shRNA knockdowns. 38/61 (62%) genes were co-ordinately differentially expressed in both analyses (Youssef et al., 2013; Figure 5A; Supplementary Table ST3).

**Figure 5.**
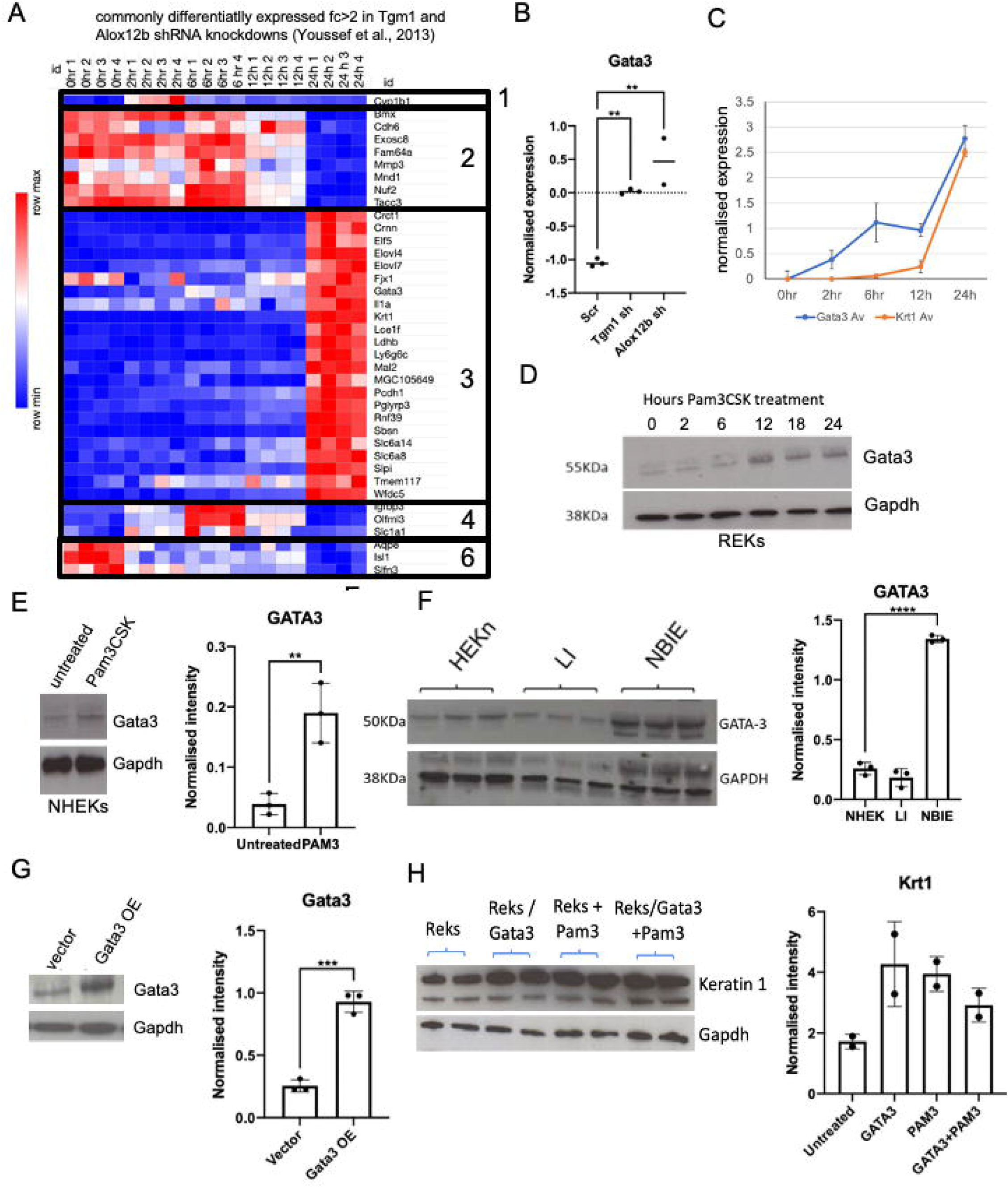
Gata3 up-regulation increased keratin 1 expression downstream of TLR2 activation. A) Heat map of genes differentially expressed in both Tgm1 and Alox12b shRNA knockdown rat epidermal keratinocytes (Youssef et al., 2013) and in the PAM3CSK4 treated keratinocytes. Differentially expressed genes are mapped to each to the Clusters identified in figure 4. B) Normalised affymetrix signals of Gata3 in Tgm1 and Alox12b shRNA knockdown and Scrambled control samples (Scr). C) Graph of normalised expression of Gata3 and Keratin 1 (Krt1) across the 24-hour time-course of PAN3CSK treatment. D) Western blot of rat epidermal keratinocytes treated with PAM3CSK4 over a 24-hour timecourse. E) Western blot of Gata3 in normal human epidermal keratinocytes (NHEKs) treated with PAM3CSK4 for 24 hours, on the right is a graphical representation of normalised densitometry data (n=3) F). Western blot of Gata3 in lamellar ichthyosis (LI) and non-bullous ichthyosiform erythroderma and normal controls. Graph shows normalised intensity. G) Western blot of Gata3 in Gata3 overexpressing cells (Gata3 OE) compared to vector alone. Graph shows normalised densitometry data (n=3). H) Western blot of duplicate rat epidermal keratinocyte cultures over-expressing Gata3, treated with PAM3CSK4 or overexpressing cells treated with PAM3CSK4 for keratin 1. Gapdh serves as loading control. Graph shows normalised intensity. Bars represent standard deviation. ** p<0.005, ***p<0.0005, ****p<0.00005, one-way ANOVA with post-hoc testing.

Gata3 was of particular interest, as it had been identified from our combined analysis of gene expression data and proteomic data. Gata3 expression was increased in both the Tgm1 and Alox12b knockdown keratinocytes, with high expression in Alox12b knockdown keratinocytes (Figure 5B). in the PAM3CSK4 treatment mRNAseq analysis, Gata3 was found in Cluster 3, the genes increased maximally at 24 hours, however Gata3 mRNA expression progressively increased during the whole PAM3CSK4 treatment period, prior to the increase in keratin 1 (Figure 5C) Gata3 protein increased after 12 hours of PAM3CSK4 treatment in rat epidermal keratinocytes (Figure 5D), and the same increase was seen in 24-hour PAM3CSK4 treated human keratinocytes (Figure 5E). We looked at the over-represented binding sites in the human promoters for the genes in Cluster 3 from ENCODE Chromatin precipitation data using Enrichr, Genes with Gata3 binding promoters were significantly over-represented in Cluster 3 genes (Supplementary figure S4A), and they were enriched for both genes involved in epidermal terminal differentiation and lipid synthesis (Supplementary figure S4B and C). This suggested that the up-regulation of epidermal differentiation genes and lipid synthesis was potentially driven by the up-regulation of Gata3.

We examined the expression of Gata3 in our 2 patient keratinocyte lines. Gata3 increased in BCIE keratinocytes but not in LI keratinocytes (Figure 5F). Overexpression of Gata3 in rat epidermal keratinocytes (Figure 5G and H) increased keratin 1 expression. However, treatment with PAM3CSK4 did not increase K1 further, suggesting that Gata3 was downstream of TLR2 activation, and potentially was a key downstream driver of the longer-term effects of TLR2 activation leading to hyperkeratosis in patients with BCIE.

## Discussion

We show that TLR signalling is activated in ARCI models, and that there is evidence for increased TLR signalling and sumoylation in scale from ARCI patients. TLR2 activation is necessary for hyperkeratosis and blockade of TLR2 signalling in ARCi models and patient keratinocyte lines reduced the increased levels keratin 1 seen in ARCI (O’Shaughnessy et al., 2010; Youssef et al., 2013). We show proof of Principal that blockade of TLR2 signalling can transiently reduce keratin 1 expression for 2 hours in patient keratinocytes. However in HEK293 cells, treatment of CU-CPT22 caused reduced nuclear NFκβ at 20 minutes (Daniele et al., 2015), while in keratinocytes inoculated *with P,acnes*, CU-CPT22 was still effective at 24 hours (Su et al., 2017). This suggests that the effects of CU-CPT22 are variable, context dependent or that the TLR2-NFκβ mediated pathway that increased keratin 1 is different, and responds differently to CU-CPT22. More investigation into both the stability of the drug and the effects of treatment in the ARCI background are required.

We found that activation of TLR2 leads to upregulation of SUMO1 in the epidermis. Typically, the activation of SUMO correlates with downregulation of the innate immune response (Hanke et al., 2014). Sumoylation increases during epidermal terminal differentiation (Deyrieux et al., 2007), and sumoylated substrate proteins were concentrated in the upper epidermis, which confers a significant proportion of skin barrier function. Inhibition of sumoylation prevented keratinocyte differentiation. This is consistent with increased sumoylation associating with a protective pro-differentiation response in individuals with ARCI. Sumolyation is known the coordinate inflammation and antiviral responses during innate sensing, with sumoylation blunting these responses in mice, due to enhancer binding in pro-inflammatory genes (Decque et al., 2016). Sumoylation caused by impaired barrier function in ARCI, may therefore be required to direct the TLR2-NFκβ response towards a barrier repair programme

Activation of TLR2 leads to Claudin upregulation (Kuo et al., 2013), and loss of TLR2 correlates with reduced barrier metrics and transepidermal water loss (Kuo et al., 2013). Therefore, it is likely that the increased activation of TLR2 signalling is a protective response to the defective barrier in ichthyosis, supporting this concept is the increased thickness of the cornified layer in organotypic cultures treated with PAM3CSK4. Activation of TLR2 by PAM3CSK4 induces upregulation of Claudin 1 and Claudin 23, and we replicated these data in our analyses. So TLR2 activation seems to rescue epidermal barrier function in part by activation of tight junctions (Yuki et al., 2011). We show by comprehensive mRNAseq analysis that the effects of TLR2 activation are far more wide-ranging than the established role in classical innate immune activation. We identified novel functions for TLR2 signalling in epidermal terminal differentiation and lipid synthesis, both of which would be required for optimal barrier restoration in response to the severe barrier defects in ARCI.

One difference between rat and human that needs to be addressed is the different TLR2 expression changes in response to PAM3CSK4 treatment, and how this may relate to differences in TLR2 expression and signalling in barrier disruption. TLR2 is constitutively expressed in human epidermis (Panzer et al., 2014), consistent with our date in keratinocytes, and protein expression levels do not change in patients with barrier dysfunction (Donetti et al., 2017). However, in mouse skin, TLR2 protein levels are increased in response to barrier disruption such as tape stripping (Jin et al., 2009). This is likely a biological difference, as it is still observed in cultured REKs and NHEKs, regardless, the downstream effect, upregulation of terminal differentiation markers remained the same. Therefore, TLR2 activation is directly linked to epidermal terminal differentiation.

One key effect of TLR2 activation by PAM3CSK4 is the biphasic activation of NFκβ signalling. This is consistent with our previous data that showed an up-regulation of nuclear p65 in Tgm1 shRNA expressing rat epidermal keratinocytes and organotypic cultures (O’Shaughnessy et al., 2010). In this manuscript we show that NFκβ related genes are markedly different in the “early” and “late” phase of the response to TLR2 activation. Our previous studies reflected the late stage of the TLR2 activation response, post 24 hours. Whilst NFκβ activation is required for the innate immune response, this was only apparent in early timepoints in our TLR2 activation study, while the second peak of NFκβ activation at the 24 hours timepoint was more clearly associated with a barrier protective response including the expression of epidermal differentiation and lipid synthesis genes. This is consistent with other work that has shown the up-regulation of NFκβ in the flaky tail mice, a model for ichthyosis vulgaris (Kypriotou et al., 2013). However, NFκβ deletion or inhibition can also result in skin hyperproliferation and inflammation (Kumari et al., 2021). Control of NFκβ signalling is complex involving both the pattern recognition receptors such as the TLRs, signalling through Myd88 and the action of IKK degrading Ικβα which allows NFkB into the nucleus (Descargues et al, 2008). Loss of IKK2 prevents epidermal terminal differentiation during murine development (Pasparakis et al., 2002). Taken together this strongly implicates nuclear translocation of NFκβ in response to TLR2 activation in activating epidermal terminal differentiation and enhancing epidermal terminal differentiation in response to the barrier dysfunction in ARCI.

GATA3 knockdown resulted in a selected barrier impairment related to the downregulation of genes involved in lipid synthesis (De Guzman Strong et al., 2006, and over-expression of GATA3 increased expression of keratinisation genes such as Loricrin, and inhibits proliferation (Kawachi et al., 2012; Masse et al., 2014; Zeitvogel et al., 2017). Up-regulation of lipid synthesis genes and epidermal terminal differentiation related genes occurred in both our PAM3CSK4 treated keratinocytes and in our Tgm1 and Alox12b shRNA knockdown keratinocytes (Youssef et al., 2013), while downregulation of cell cycling occurred in response to PAM3CSK4 treatment specifically in the Alox12b shRNA knockdown keratinocytes (Youssef et al.,2013). GATA3 plays an important role in aberrant epidermal terminal differentiation. Methylation of GATA3 by the drug DZ2002 reduces psoriasis skin lesions (Chen et al., 2021). There is evidence to suggest GATA-3 is a target gene of p63 and is involved in upregulation of IKKα, also involved in epidermal differentiation (Candi et al, 2006), linking GATA3 up-regulation to the NFKB response in our Cluster 3 genes. We therefore have defined a role for TLR2, NFκβ and GATA3 in the establishment of altered epidermal differentiation programmes in response to diseases of epidermal barrier function, and in particular the hyperkeratotic program established in response to barrier dysfunction in ARCI. One important outstanding question that may have further therapeutic implications is that TLR2 signalling in our ARCI shRNA models is increased, suggesting that there is an endogenous ligand that is present in defective differentiated cells that is released into the medium. Although outside of the scope of this manuscript, Heat shock proteins HSP60 and HSP70 are abundant in differentiating keratinocytes and is known to activate TLR2 signalling (Vabulus et al., 2002, Habich et al., 2002). Understanding the extracellular milieu in barrier defective skin may reveal further therapeutic targets to reduce hyperkeratosis induced by TLR2 activation.

## Acknowledgements

This work was supported by Grant funding from the British Skin Foundation, Foundation for Ichthyosis and Related Skin Types (F.I.R.S.T.), and Great Ormond Street Childrens’ Charity. We thank the QMUL Genome Centre for the mRNAseq and subsequent analysis. RFLO’S, JIH and KM conceived the study. RFLO’S, HT and SH designed experiments. HT, SH, and WH performed experiments and interpreted data. The manuscript was written by RFLO’S, KM and HT and was contributed to by all authors.

## Supplementary information

**Supplementary figure S1: PAM3CSK44 upregulates Sumo1, Keratin 1 and IL1**α **but not keratin 10**. B) Western blot analysis of keratin 1, keratin 10 and IL1α in rat epidermal keratinocytes treated with various stress inducers compared to untreated controls. GAPDH served as loading control.C) Normalised densitometry of western blots untreated controls. *p = <0.05, **p = < 0.005, One-way ANOVA with post-hoc testing.

**Supplementary figure S2: Tlr2 mRNA expression not altered in ARCI shRNA expressing models**. Graphs shows normalised expression of Tlr family genes (n=3, Scram; n=3 Tgm1 shRNA, n=2 Alox12b shRNA). ***, p<0,0005 ANOVA with post-hoc testing.

**Supplementary figure S3: Tlr expression in response to PAM3CSK4 treatment**. Graphs show RPKM values for the Tlrs shown across 24 hours of PAM3CSK4 treatment. *, p <0.05, ** p<0.005, ***p<0.0005, ****p<0.00005, one-way ANOVA with post-hoc testing.

**Supplementary figure S4: GATA3 binds promoters of Cluster 3 genes involved specifically in lipid metabolism and epidermal differentiation**. A) Cluster 3 genes were interrogated with Enricher to find GATA3 binding promoters in the human homologue of the gene. B) Graph of over-represented gene ontologies associated with the subset of genes bound by GATA3 in cluster 3. C) STRING network genes of these proteins showed a large, interrelated cluster incorporating genes involved in lipid metabolism and epidermal development.

**Supplementary table ST1: Summarised mass spectrometry data**. Normalised protein abundance for cornified envelopes from 8 normal controls (N1-8) and 4 ARCI patients (LI1-4), showing average fold changes (FC) and uncorrected p-values (p-Val) of T-tests across the 4 patients. Red and green – increased and decreased expression respectively.

**Supplementary table ST2: Normalised expression data across the 24-hour timecourse of PAM3CSK4 treatment and K means clustering of the dataset**. 1-6 represent the clusters identified and shown on figure 4, n=4 each timepoint

**Supplementary table ST3: Normalised expression of all genes altered by PAM3CSK4 treatment and differentially expressed in both Tgm1 and Alox12b shRNA knockdowns (Youssef et al., 2013)**. 1-6 represent the clusters identified and shown on figure 5, n=4 each timepoint.

**Supplementary table ST4: RPKM-normalised data for all transcripts in at all timepoints of PAM3CSK4 treatment**. N=4 each timepoint.

