## Supplementary Information for "Chronic activation of Toll-like receptor 2 induces an ichthyotic skin phenotype"

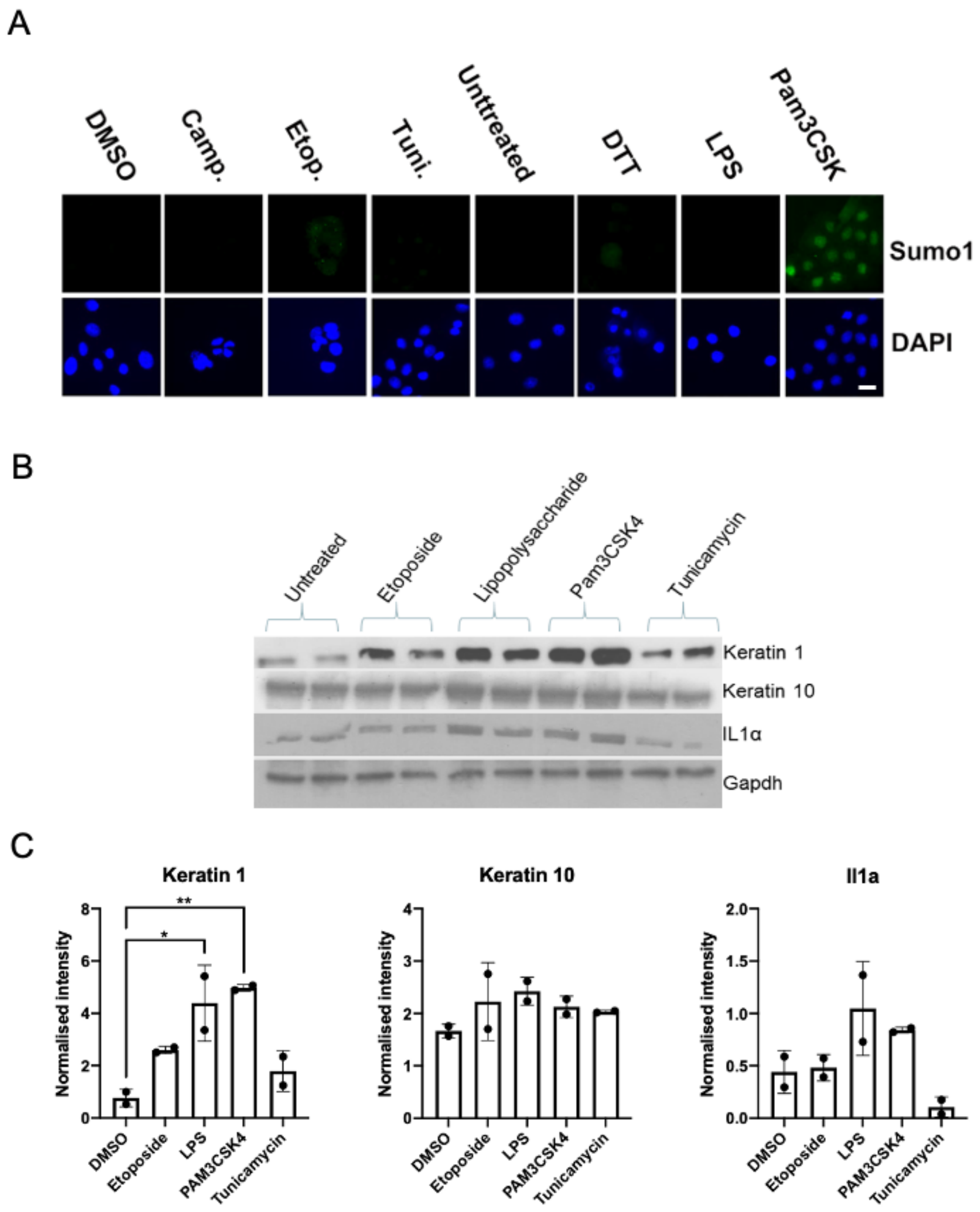


Supplementary figure S1(Revised)

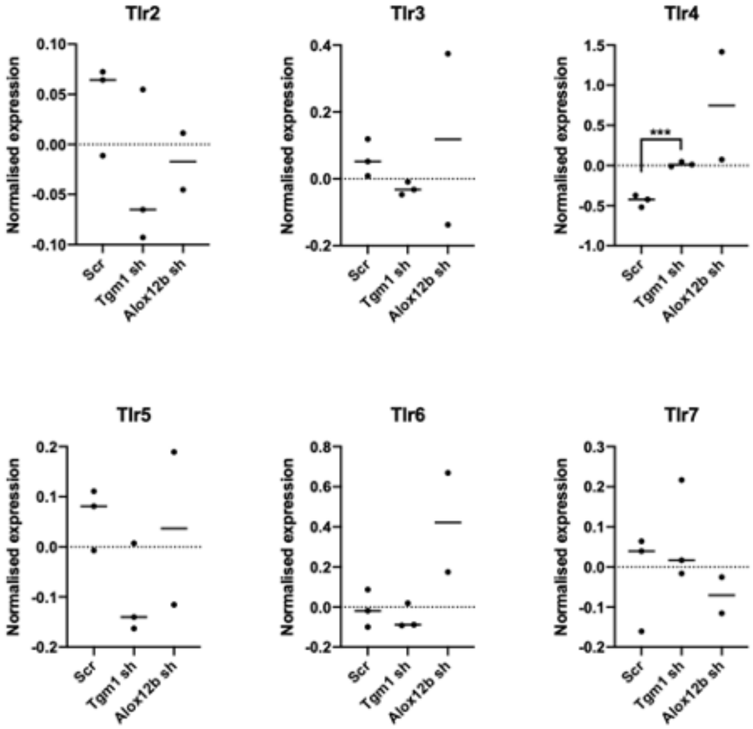


Supplementary figure S2 (Revised)

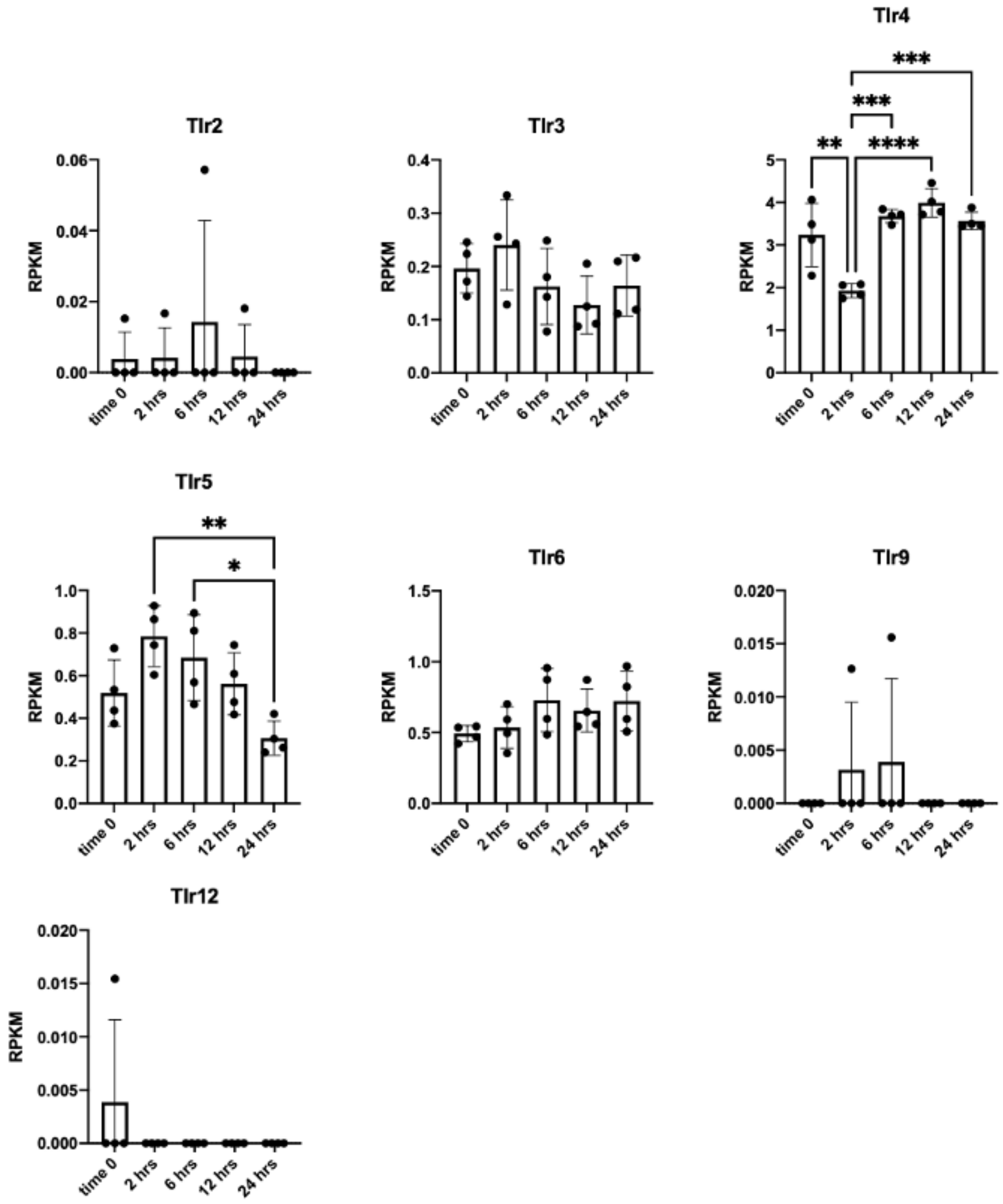


Supplementary figure S3 (Revised)

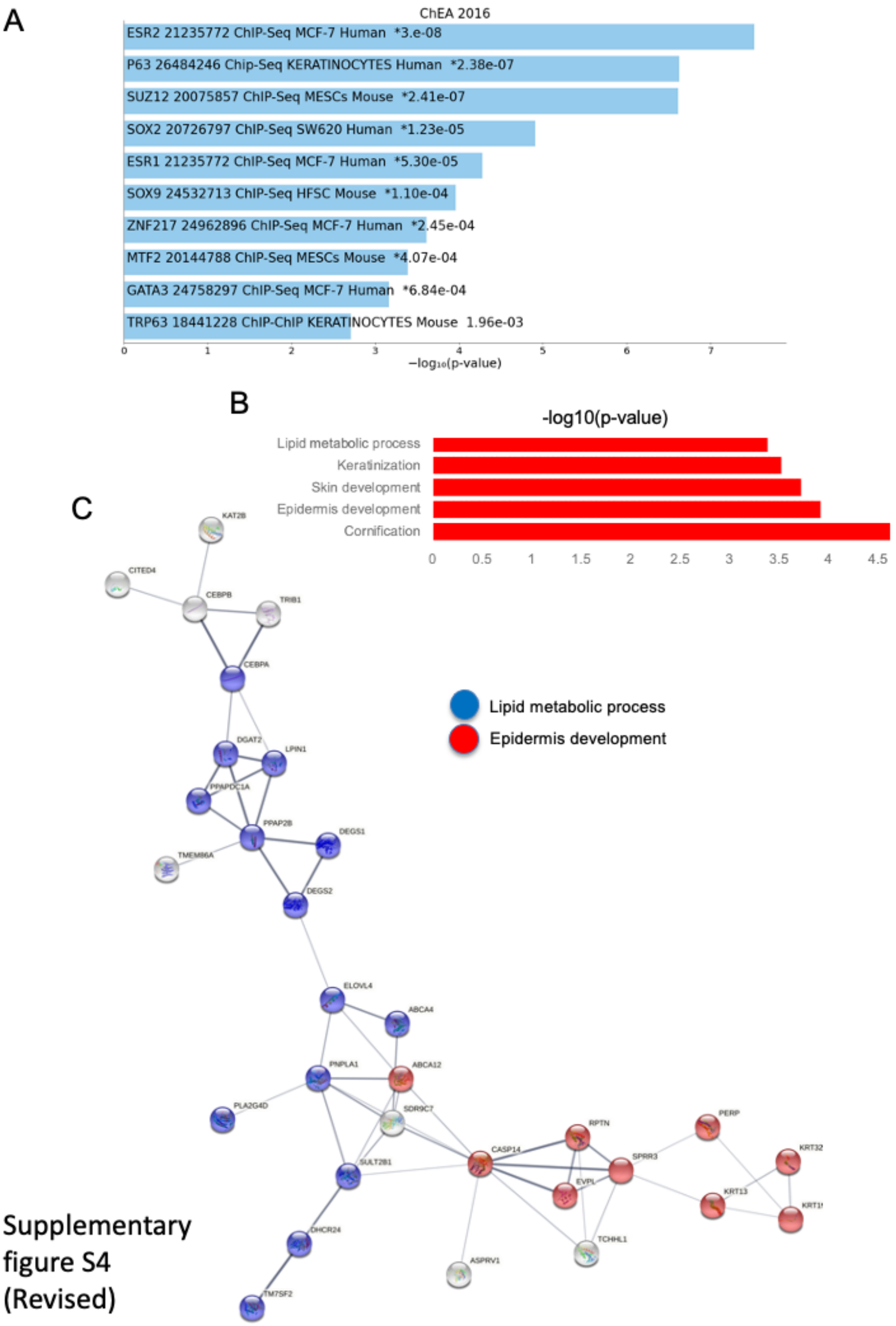


Supplementary figure S4 (Revised)
